# Three-dimensional chromatin landscapes in MLLr AML

**DOI:** 10.1101/2023.11.02.565383

**Authors:** Pinpin Sui, Peng Zhang, Feng Pan

## Abstract

Rearrangements of the mixed lineage leukemia (MLLr) gene are frequently associated with aggressive acute myeloid leukemia (AML). However, the treatment options are limited due to the genomic complexity and dynamics of 3D structure, which regulate oncogene transcription and leukemia development. Here, we carried out an integrative analysis of 3D genome structure, chromatin accessibility, and gene expression in gene-edited MLL-AF9 AML samples. Our data revealed profound MLLr-specific alterations of chromatin accessibility, A/B compartments, topologically associating domains (TAD), and chromatin loops in AML. The local 3D configuration of the AML genome was rewired specifically at loci associated with AML-specific gene expression. Together, our results show that the 3D chromatin undergoes extensive reorganization at multiple architectural levels, which underpins the remodeling of the transcriptome in MLLr AML.

## Results

The clinically important and genetically well-defined MLLr leukemia accounts for up to 50% of infant and 10% of adult acute leukemia that are associated with very poor prognosis and chemo-resistance [1-3]. The current mechanistic understanding of MLLr AML initiation and progression have not yet translated into therapeutic success due to the lack of an accurate and reliable human model, and the complexity of genomic events contributing to disease. Recent CRISPR/Cas9-mediated generation of MLLr in human hematopoietic stem and progenitor cells (HSPCs) has enabled the modeling of key aspects of AML biology [4, 5]. To understand how the 3D topology of the genome contributes to the leukemogenesis of MLLr AML, we coupled assay for transposable-accessible chromatin using sequencing (ATAC-seq) and RNA sequencing (RNA-seq) to enhanced high-resolution chromosome conformation capture (Micro-C) in gene-edited MLL-AF9 AML cells (Fig. 1A). Normal human umbilical cord blood CD34+ cells were selected as controls since they are considered to be healthy donor cells of origin for AML. We firstly estimated the similarity between AML samples and HSPCs using principal component analysis (PCA) and differential chromatin accessibility analysis of ATAC-seq profiles (Fig. 1B-C). AML-specific accessible DNA regions exhibited a significant AML signature, indicating that these cells are a suitable experimental system to map the 3D genome architecture of AML (Fig. S1A-D). On average, around 800□million paired-end reads were generated in each Micro-C library. Unsupervised hierarchical clustering using Micro-C matrices clearly separated HSPCs and leukemia cells (Fig. S2A), indicating a specific chromatin structural landscape of MLLr AML. Notably, there were substantial changes in compartmentalization in leukemia cells vs in HSPCs (Fig. 1D-E and Fig. S2B). About 1,476 A-to-B compartment switch and 2,251 B-to-A switch were observed when comparing AML samples with HSPCs at 100 kb resolution (Fig. 1D). Genes in the A-to-B or B-to-A switching regions had decreased or increased expression, depending on the direction of the switch (Fig. 1F). To further explore the chromatin structure of MLL-AF9 AML, we identified the difference of TAD and the insulation score at TAD boundary between AML cells and HSPCs at 25 kb resolution (Fig. 1G-I and Fig. S2C-F). Correlation between AML and HSPC TAD boundaries demonstrated that over 90% TADs were shared (Fig. 1G). However, MLL translocation resulted in a significant loss of TAD strength, whereas this translocation depressed the insulation between neighboring TADs (Fig. 1H-I).

**Figure 1:**
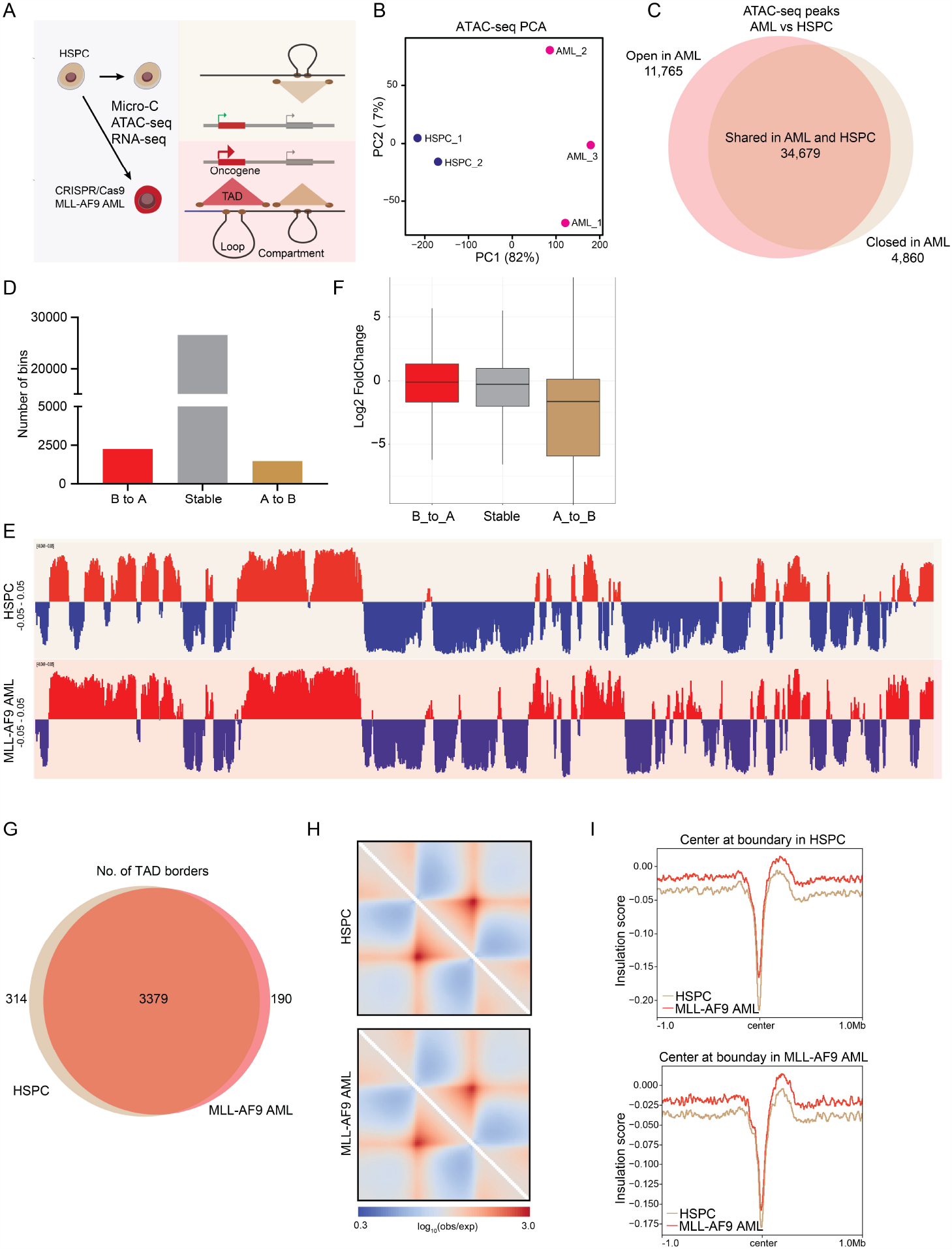
Large-scale 3D genome organization in MLL-AF9 AML. (A) Schematic overview of the 3D genome characterization of MLL-AF9 AML cells generated through CRISPR/Cas9-mediated gene editing. (B) PCA of chromatin accessibility profiles of HSPCs and gene-edited MLL-AF9 AML cells. (C) Venn diagram representation of the differential chromatin accessibility between ATAC-seq peaks of HSPC and MLL-AF9 AML. (D) Number of A/B□compartment switching in MLL-AF9 AML samples compared with HSPCs. (E) PCA analysis for the first principal component representing compartment A and B at 100 kb resolution on chr13. (F) Gene expression alterations associated with changes in the A/B□compartment. (G) Overlap of HSPC TADs and MLL-AF9 AML TADs. (H) Aggregate domain analysis (ADA) showing all TAD in HSPC (top) and MLL-AF9 AML (bottom). (I) Mean plot describes genome-wide insulation score around HSPC and MLL-AF9 AML TAD boundaries. The X-axis represents the genome distance to the TADs boundaries, while the Y-axis the insulation score.

Chromatin loops were detected using Mustache at 10 kb resolution [6]. This identified 3,866 distinct healthy donor-specific loops and 2,360 AML-specific loops (Fig. 2A). The MLL-AF9 AML-specific loops also showed subtype-specific patterns and contained many known MLL target genes, such as *UBE2J1, PARP8*, and *PHC2*, and non-coding elements in the loop anchors (Fig. 2B-D and Fig. S3A-B). Similarly, the HSPC-specific loops contained HSPC signature genes (*KIT, NRIP1*, and *IGF1R*). The MLL-AF9 AML-associated loops demonstrated enrichment for several key transcription factor (TF) motifs suggesting an interactive TF network in MLL-AF9 AML (Fig. 2E). The motifs constituted binding sites for TFs involved in hematopoietic lineage specification (GATA3) and MLL leukemia pathogenesis (MEIS1 and MYB). Recent studies indicate that chromatin loops that connect promoters to enhancers and silencers are essential for gene regulation [7, 8]. In support of this notion, we noticed 597 genes were significantly upregulated and 1368 genes were downregulated in dysregulated AML loops (Fig. S3C-F).

**Figure 2:**
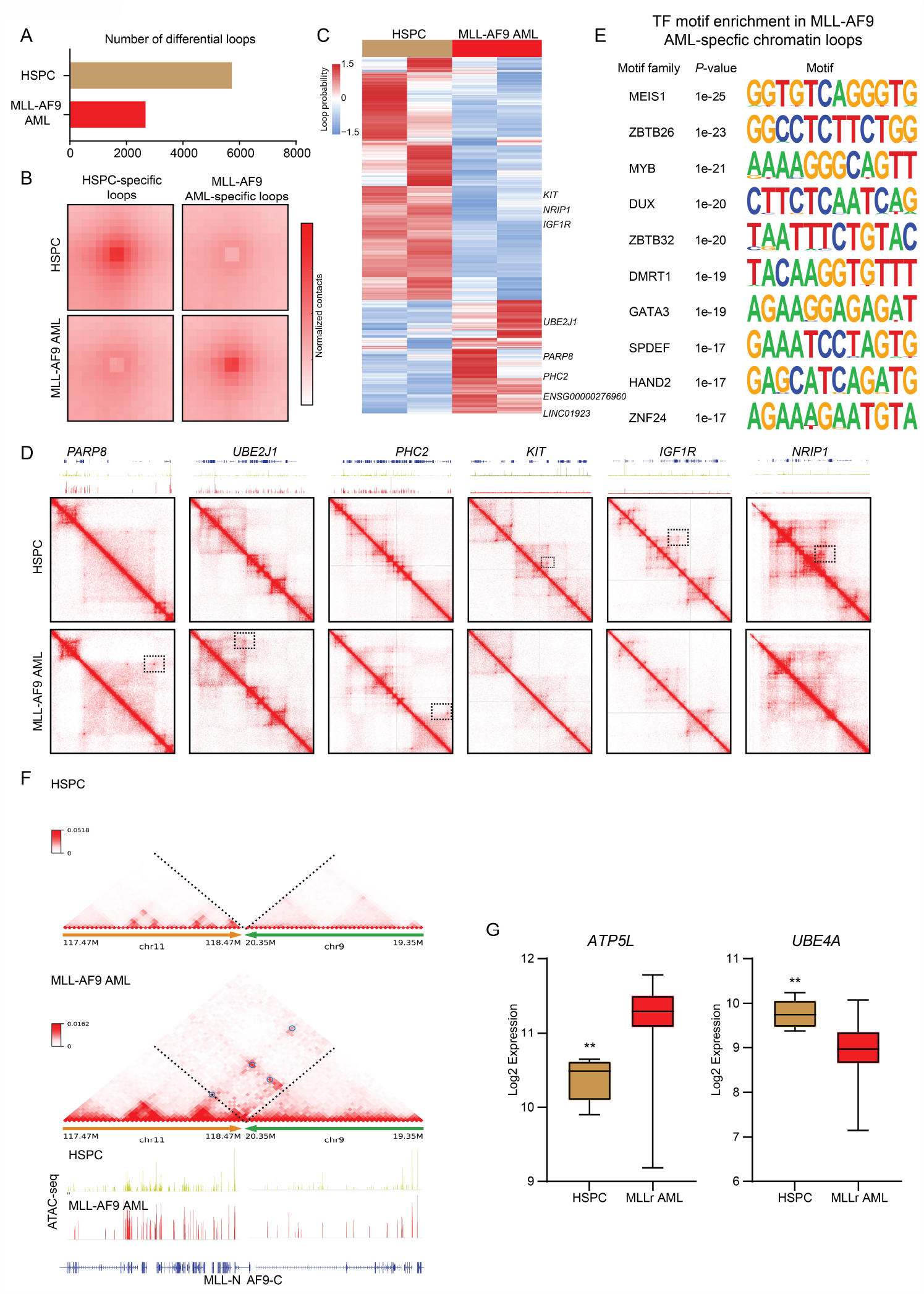
MLL-AF9 AML specific chromatin loops. (A) Number of HSPC- and MLL-AF9 AML-specific loops using the mustache at 10 kb resolution. (B) APA plot for MLL-AF9 AML interaction-increased and interaction-decreased loops compared with HSPC. (C) Heatmap shows the subtype-specific loop analysis for HSPCs and MLL-AF9 AML samples. Each row is a loop and the values are the normalized interaction frequency extracted by Juicer (Observed/Expected). (D) Micro-C interaction matrix and chromatin accessibility surrounding HSPC- and MLL-AF9 AML-specific genes. (E) De novo motif analysis showing the enrichment of transcription factors in the anchor regions of MLL-AF9 AML-specific chromatin loops using HOMER. (F) Enhancer-hijacking and silencer-hijacking events analyzed by NeoLoopFinder (black circles). (G) Box plot of ATP5L and UBE4A expression levels in normal HSPCs and MLLr AML cells from BloodSpot [10].

The observed AML bearing MLL-AF9 translocation raised the possibility that the structural variation can disrupt three-dimensional genome organization and induce enhancer/silencer hijacking. To assess potential chromatin interactions induced by translocation, the NeoLoopFinder tool was used to identify genes associated with enhancer/silencer hijacking [9]. A representative example is shown in Fig. 2F, which shows the fusion between chromosome□11 and chromosome□9 in AML but not in HSPC. This neo-loop connected *ATP5L* (located on chromosome□11) to several enhancers on chromosome□9. By contrast, no such inter-chromosomal Micro-C signals were observed in HSPC. As expected, *ATP5L* showed increased expression in MLLr AML samples that exhibited enhancer hijacking. However, *UBE4A* showed decreased expression in MLLr AML samples, suggesting silencer-hijacking events were identified in MLLr AML samples (Fig. 2G).

Together, this study presents the first three-dimensional chromatin landscapes in MLLr AML, with limitations of small sample size and lack of validation of identified loops. An interesting future experiment would be to incorporate ChIP-seq for CTCF, H3K27ac, and H3K27me3 to systematically separate loop-associated enhancers and silencers.

## Availability of data and materials

The raw data of RNA-Seq and Micro-C reported in this paper have been deposited in the Gene Expression Omnibus database (accession number GSE244472).

## Acknowledgements

The authors thank Michael Cleary for generously providing gene-edited cells, and Mingjiang Xu for constructive discussions and comments, and Shi Chen, Juan Wang for animal assistance. This work was supported by funding from the Alex’s Lemonade Stand Foundation and UT Health San Antonio Mays Cancer Center Early Career Grant to F.P.

## Authorship contributions

PS and FP conceived the project and wrote the manuscript. PS, PZ, and FP designed and performed experiments, and analyzed data. All authors read and approved the final manuscript.

