## Supplementary data for "Three-dimensional chromatin landscapes in MLLr AML"

**Materials and methods**

**Cell culture**

Human CD34$^+$ HSPCs and CRISPR-Cas9 engineered MLL-AF9 leukemia cells were cultured in StemSpan SFEM II medium with SCF (50 ng/mL), thrombopoietin (100 ng/mL), Flt3 ligand (100 ng/mL), IL-6 (100 ng/mL), IL-3 (50 ng/mL), G-CSF (50 ng/mL), UM729 (0.75 µM), StemRegenin 1 (0.75 µM), and 20% FBS at 37°C, 5% $CO_2$ as previously described [1].

**Micro-C library preparation and data analysis**

Micro-C libraries were performed following Dovetail Micro-C Kit protocol (Cat# 21006, Dovetail Genomics, Scotts Valley, CA, USA). Briefly, 1x10$^6$ leukemia and healthy donor cells were resuspended in 1xPBS and crosslinked with 3mM DSG and 1% formaldehyde. Treated cells were then washed and digested with 0.5 uL MNase in 100 uL of Nuclease digest buffer. Digested chromatin was thawed by addition of 3µl of 20% SDS and capture on chromatin capture beads. Chromatin bound DNA fragments were end repaired and A-tailed using the End Polishing enzyme mix and buffer. Dovetail bridge adapters were ligated to the free ends of chromatin-bound DNA fragments, following the Micro-C kit protocol. This was followed by intra-aggregate ligation to form chromatin-chromatin long-range interactions. Finally, the chromatin-bound DNA was recovered by treating the sample with proteinase K to reverse the crosslinks, and then purifying the DNA using SPRIselect beads (Beckman Coulter). After DNA purification, PCR was performed to amplify the library and size range between 350–1,000 bp was selected for the library. The final libraries were sequenced by Illumina NovaSeq 6000 with the mode of paired-end 150bp. Trim galore (v 0.6.7) (https://github.com/FelixKrueger/TrimGalore) was used to trim the raw sequencing data with the parameter of –paired -q 25 –phred33 –length 35 -e 0.1 –stringency 2. Clean reads from each library were aligned to the human genome (GRCh38) using BWA (v 0.7.17-r1188) [2]. Pairtools (v1.0.2) was used to process the alignment output and generate the final bam file used for the contact matrix [3]. Juicer (v 1.22.01) and cooler (v 0.9.1) were used to generate .cool and .hic file respectively [4, 5]. A/B compartments were identified by FAN-C (v

0.9.25) at 100 kb resolution [6]. TADs were identified by juicer arrowhead (v 1.22.01) at 25 kb resolution. Loops were identified by mustache (v 1.3.1) at 10 kb resolution [7]. Insulation scores were calculated by cooltools (v 0.5.4) at 25 kb resolution (https://doi.org/10.1101/2022.10.31.514564). Aggregate domain analysis (ADA) was performed with coolpup.py (v 1.1.0) at 25 kb resolution [8].

**ATAC-seq analysis**

ATAC-seq was performed as described [9, 10]. Briefly, healthy donor and AML cells were lysed in cold lysis buffer (10 mM Tris-HCl, pH 7.4, 10 mM NaCl, 3 mM MgCl2, and 0.1% IGEPAL CA-630). Nuclei were centrifuged at 500 × g for 10 min, 4°C. Nuclei extract were then incubated with Nextera Tn5 Transposase, 2× TD buffer, and nuclease free water at 37°C for 30 min with gentle mixing. After DNA purification with the MinElute PCR Purification Kit (Qiagen), PCR was performed to amplify the library for additional cycles distinctively according to a quantitative PCR reaction for optimum cycles. The final libraries were sequenced by Novogene. Adapter sequences were trimmed and reads were mapped to Hg38 using Bowtie2 [11]. ATAC-seq peak calling was performed with Genrich (https://github.com/jsh58/Genrich) with default setting.

**RNA-seq analysis**

RNA was extracted from control and leukemia cells with RNeasy Plus Mini Kit (Qiagen) according to the manufacturer's instructions. RNA-seq libraries were generated and sequenced by Illumina NovaSeq 6000 (2x150bp). Trim galore (v 0.6.7) was used to trim the raw data with the parameter of –paired -q 25 –phred33 –length 35 -e 0.1 –stringency 2 (https://github.com/FelixKrueger/TrimGalore). The clean reads were aligned to the human genome (GRCh38) using STAR (v 2.7.10a) [12] and then HTSeq (v 2.0.2) [13] was used to calculate the raw read counts for each gene and each sample. The count matrix was used to identify differentially expressed genes (DEGs) by DESeq2 (v 1.36.0) [14] with the cutoff of FDR < 0.05 and |Fold change| >2.

Figure S1

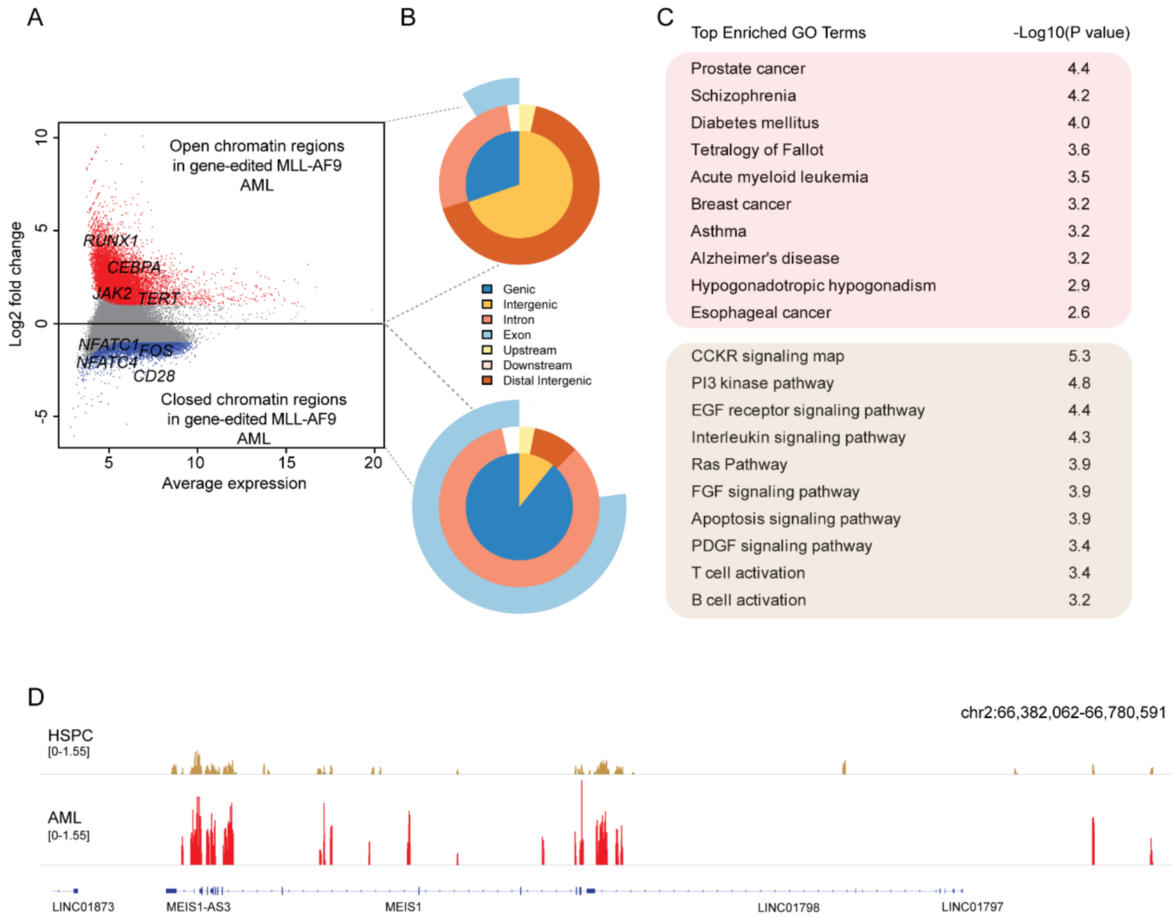
A
Open chromatin regions in gene-edited MLL-AF9 AML
RUNX1
CEBPA
JAK2
TERT
NFATC1
FOS
NFATC4
CD28
Closed chromatin regions in gene-edited MLL-AF9 AML
Log2 fold change
Average expression
B
Genic
Intergenic
Intron
Exon
Upstream
Downstream
Distal Intergenic
C
Top Enriched GO Terms
-Log10(P value)
Prostate cancer 4.4
Schizophrenia 4.2
Diabetes mellitus 4.0
Tetralogy of Fallot 3.6
Acute myeloid leukemia 3.5
Breast cancer 3.2
Asthma 3.2
Alzheimer's disease 3.2
Hypogonadotropic hypogonadism 2.9
Esophageal cancer 2.6
CCKR signaling map 5.3
PI3 kinase pathway 4.8
EGF receptor signaling pathway 4.4
Interleukin signaling pathway 4.3
Ras Pathway 3.9
FGF signaling pathway 3.9
Apoptosis signaling pathway 3.9
PDGF signaling pathway 3.4
T cell activation 3.4
B cell activation 3.2
D
chr2:66,382,062-66,780,591
HSPC
[0-1.55]
AML
[0-1.55]
LINC01873
MEIS1-AS3
MEIS1
LINC01798
LINC01797

**Figure S1: Comparison of chromatin accessibility between HSPC and MLL-AF9 AML.**

(A) MA plot of the normalized ATAC-seq intensities of cell type-specific peaks from healthy donor and diseased individuals.

(B) Distribution of genomic features of differential regulatory elements.

(C) Top enriched Gene Ontology terms of differential peaks.

(D) Normalized ATAC-seq signal profiles at *MEIS1* locus in HSPC and MLL-AF9 AML.

Figure S2

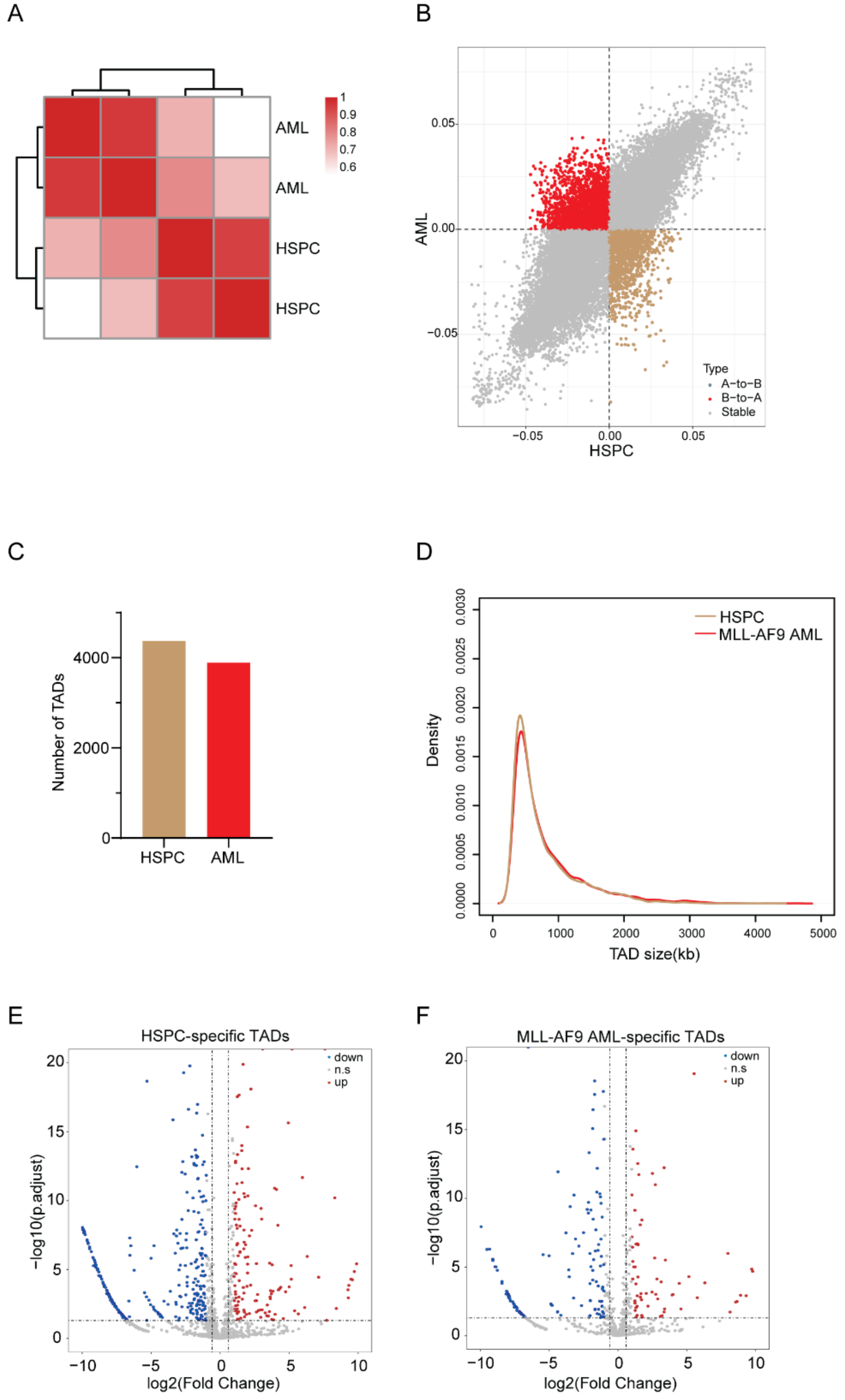

A
AML
AML
HSPC
HSPC
1
0.9
0.8
0.7
0.6
B
AML
HSPC
0.05
0.00
−0.05
−0.05
0.00
0.05
Type
A−to−B
B−to−A
Stable
C
Number of TADs
4000
2000
0
HSPC
AML
D
Density
0.0000
0.0005
0.0010
0.0015
0.0020
0.0025
0.0030
HSPC
MLL-AF9 AML
0
1000
2000
3000
4000
5000
TAD size(kb)
E
HSPC-specific TADs
−log10(p.adjust)
log2(Fold Change)
down
n.s
up
F
MLL-AF9 AML-specific TADs
−log10(p.adjust)
log2(Fold Change)
down
n.s
up

**Figure S2: A/B compartment switch and TAD alterations between HSPC and MLL-AF9 AML.**

(A) Unsupervised hierarchical clustering of the Micro-C signals between all the samples. Each row and each column is one sample.

(B) Dot plot of eigenvector shows the A/B compartment switch among HSPC and MLL-AF9 AML.

(C) Number of TADs in HSPC and MLL-AF9 AML.

(D) Size distribution of TADs of HSPC and MLL-AF9 AML at 25 kb resolution.

(E) Differential expression analysis of genes located inside HSPC-specific TADs.

(F) Differential expression analysis of genes located inside MLL-AF9 AML-specific TADs.

Figure S3

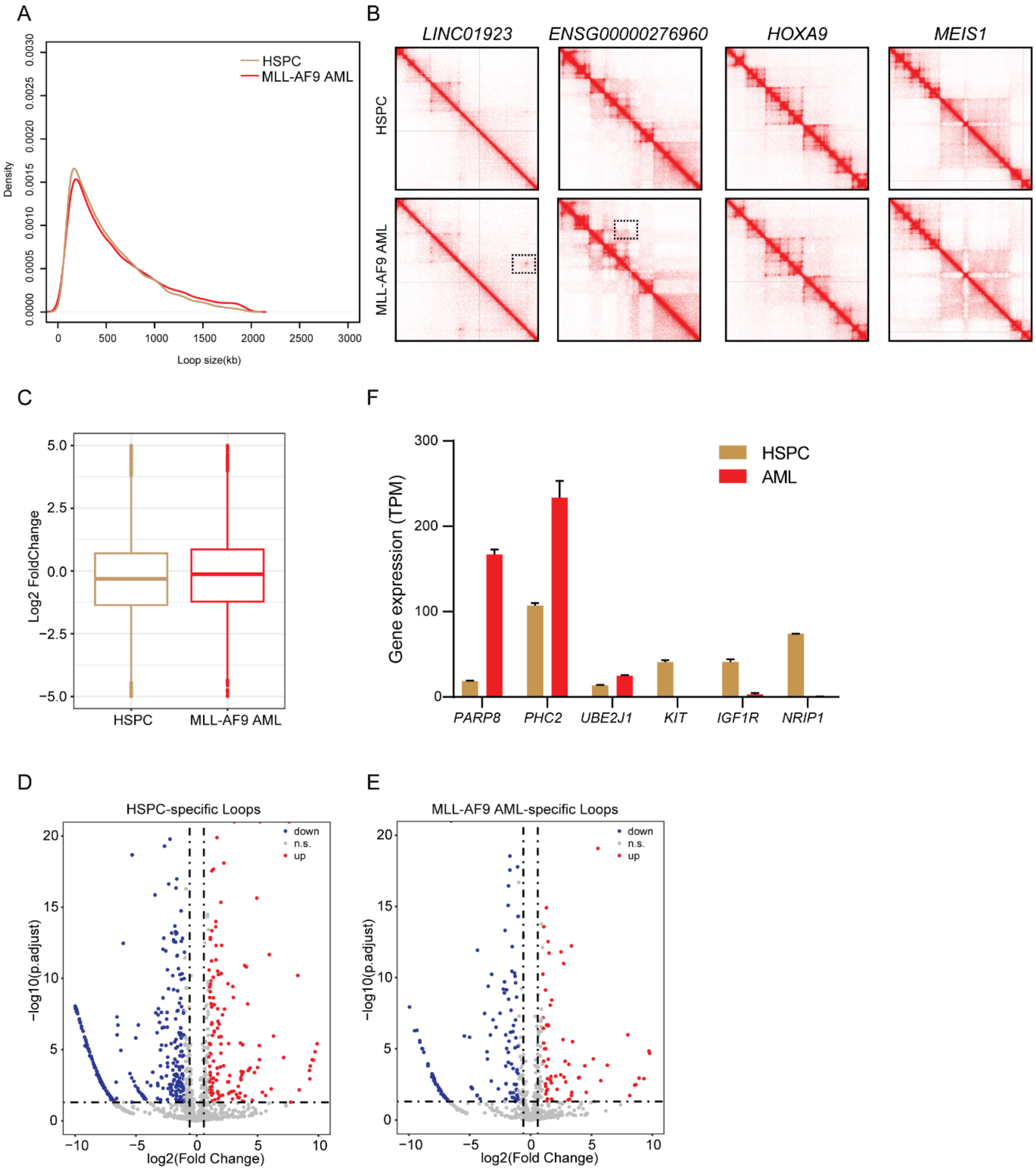
A
HSPC
MLL-AF9 AML
Density
Loop size(kb)
B
LINC01923
ENSG00000276960
HOXA9
MEIS1
HSPC
MLL-AF9 AML
C
Log2 FoldChange
HSPC
MLL-AF9 AML
F
Gene expression (TPM)
HSPC
AML
PARP8
PHC2
UBE2J1
KIT
IGF1R
NRIP1
D
HSPC-specific Loops
down
n.s.
up
−log10(p.adjust)
log2(Fold Change)
E
MLL-AF9 AML-specific Loops
down
n.s.
up
−log10(p.adjust)
log2(Fold Change)

**Figure S3: MLL-AF9 AML-specific loop analysis.**

(A) Size distribution of loops of HSPC and MLL-AF9 AML at 10 kb resolution.

(B) Micro-C matrix surrounding AML noncoding elements (LINC01923 and ENSG00000276960) and MLLr signature genes (*HOXA9* and *MEIS1*).

(C) Gene expression alterations associated with loop changes.

(D) Differential expression analysis for genes in HSPC-specific loops.

(E) Differential expression analysis for genes in MLL-AF9 AML-specific loops.

(F) The gene expression (TPM) of HSPC and MLL-AF9 AML specific genes.
